# Optimized high-throughput screening of non-coding variants identified from genome-wide association studies

**DOI:** 10.1101/2022.03.11.483985

**Authors:** Tunc Morova, Yi Ding, Chia-Chi Flora Huang, Funda Sar, Tommer Schwarz, Claudia Giambartolomei, Sylvan C. Baca, Dennis Grishin, Faraz Hach, Alexander Gusev, Matthew L. Freedman, Bogdan Pasaniuc, Nathan A. Lack

**Affiliations:** Vancouver Prostate Centre, Vancouver, BC V6H 3Z6, Canada; Bioinformatics Interdepartmental Program, University of California, Los Angeles, Los Angeles, CA 90095, USA; Central RNA Lab, Istituto Italiano di Tecnologia, Genova 16163, Italy; Department of Pathology and Laboratory Medicine, David Geffen School of Medicine, University of California, Los Angeles, Los Angeles, CA 90095, USA; Department of Medical Oncology, The Center for Functional Cancer Epigenetics, Dana Farber Cancer Institute, Boston, MA 02215, USA; Department of Urologic Science, University of British Columbia, Vancouver, BC V5Z 1M9, Canada; Department of Epidemiology, Harvard T.H. Chan School of Public Health, Boston, MA 02115, USA; The Center for Cancer Genome Discovery, Dana Farber Cancer Institute, Boston, MA 02215, USA; Department of Human Genetics, David Geffen School of Medicine, University of California, Los Angeles, Los Angeles, CA 90095, USA; Department of Computational Medicine, University of California, Los Angeles, Los Angeles, CA 90095, USA; School of Medicine, Koç University, Istanbul, 34450, Turkey

## Abstract

The vast majority of disease-associated single nucleotide polymorphisms identified from genome-wide association study (GWAS) are localized in non-coding regions. A significant fraction of these variants impact transcription factors binding to enhancer elements and alter gene expression. To functionally interrogate the activity of such variants we developed snpSTARRseq, a high-throughput experimental method that can interrogate the functional impact of hundreds to thousands of non-coding variants on enhancer activity. snpSTARRseq dramatically improves signal-to-noise by utilizing a novel sequencing and bioinformatic approach that increases both insert size and number of variants tested per loci. Using this strategy, we interrogated 70 of 140 known prostate cancer (PCa) risk-associated loci and demonstrated that 26 (37%) of them harbor 36 SNPs that significantly altered enhancer activity. Combining these results with chromosomal looping data we could identify interacting genes and provide a mechanism of action for 20 PCa GWAS risk regions. When benchmarked to orthogonal methods, snpSTARRseq showed a strong correlation with *in vivo* experimental allelic-imbalance studies whereas there was no correlation with predictive *in silico* approaches. Overall, snpSTARRseq provides an integrated experimental and computational framework to functionally test non-coding genetic variants.

## Introduction

Germline genetic variants contribute to numerous diseases from COVID-19 (1) to cancer development (2–6). Disease associated SNPs are primarily identified from genome wide-association studies (GWAS) (7). While those SNPs that occur in protein-coding regions have a predictable impact on protein sequence, the vast majority of disease-associated SNPs are located in non-coding regions (8, 9). There is increasing evidence that these non-coding variants affect disease initiation and progression by altering critical *cis-*regulatory elements (CRE) that are involved in the spatiotemporal expression of target genes (10–14). These variants commonly occur at enhancers where they can alter transcription factors (TF) binding and gene transcription (15–19). For instance, a SNP in the PCAT19 locus disrupts NKX3.1 and YY1 binding which alters enhancer activity causing dysregulation of oncogene expression and prostate cancer (PCa) progression (14, 20, 21). While most SNPs identified from GWAS studies are found within the non-coding region of the genome, it remains difficult to mechanistically characterize the impact of these variants (22, 23).

Several *in silico* and experimental approaches are commonly used to characterize potential pathogenic non-coding variants. Current *in silico* methods apply machine learning that is trained on previously published data to predict activity without experimental input (23–26). Given their relative ease, these bioinformatic approaches are commonly used to stratify candidates for validation. Yet, they do have several major limitations. Previous benchmarking studies demonstrated considerable variability between each computational method (27) and a very high error rate (28, 29). In contrast, experimental approaches are much more robust. One promising method utilizes *in vivo* chromatin immunoprecipitation with sequencing (ChIPseq) to quantify the impact of non-coding variants on allelic-imbalance of TF binding (30–34). Demonstrating the potential utility, recent work combined >7000 ChIPseq from 649 cell lines and identified 270,000 SNPs with altered TF binding affinity (25). While promising, this experimental approach is limited by the frequency of each variant in the tested population which requires an extremely large number of clinical samples for accurate functional genomic testing. These challenges therefore limit allelic-imbalance studies to only a handful of specific experimental models, tissues and disease states (35). To overcome these challenges, massively parallel reporter assay (MPRA) (36) and self-transcribing active regulatory region sequencing (STARRseq) (37) have been used to directly quantify the activity of thousands of non-coding sequences in a single experiment (38–41). Importantly, these methodologies do not require clinical samples. Several studies have adopted STARRseq to systematically screen the impact of SNPs on enhancer function (42–44). While this work demonstrates the feasibility of large-scale functional screens, they have been limited by poor signal-to-noise, false positives or limited statistical robustness. Further, it is unclear how these plasmid-based methods compare to *in silico* predictions and other experimental approaches.

In this work we developed a standardized experimental and computational STARRseq framework to identify those disease associated genetic variants that impact enhancer activity. This can accurately identify critical non-coding SNPs and test hundreds to thousands of variants in a single experiment. Using this approach we functionally tested 70 SNPs from known prostate cancer (PCa) risk-associated loci and demonstrated that 36 (51%) of them significantly altered enhancer activity. Combining these results with chromosomal looping we provided a mechanism of action for 20 PCa GWAS risk regions. Our methodology, snpSTARRseq, provides streamlined bioinformatic analysis and is amenable to different genomic regions, diseases and sequencing approaches. Overall, snpSTARRseq functionally characterizes hits from GWAS studies and provides mechanistic understanding of critical genetic variants.

## Results

### Design considerations and enhancements

To develop a methodology that functionally characterizes the impact of genetic variants on enhancer activity we adopted STARRseq, given both the ease of use and demonstrated experimental feasibility (42–44). We designed our experimental approach with the following features: (1) to scale efficiently allowing hundreds or thousands of variants to be tested, (2) to maintain a large insert size to ensure a high enhancer signal, (3) to have a high experimental signal to noise ratio by increasing the number of tested plasmids, (4) to reduce systemic false-positives associated with the earlier STARR plasmids. With this framework we optimized several key parameters with following improvements. To reduce false-positives we utilized the second-generation STARRseq plasmid and ensured that the experimental models had minimal IFN-gamma response, which can strongly influence STARRseq signal (45). Next, we utilized a DNA-capture to enrich the number of plasmids per variant and increase the signal to noise. With a low number of plasmids per variant the results are extremely error prone as you cannot separate the impact of insert size from genetic variant (45, 46). Further, the use of DNA capture also eliminated the fragment size limitations that are intrinsic with oligonucleotide synthesis (200-300bp) (47, 48). Increasing the library insert size leads to an increase in relative signal strength which reduces false positives and negatives (46, 49). We also incorporated a unique molecular identifier (UMI) for each cloned fragment to increase statistical strength and limit amplification artifacts. Finally, we used pooled genomic material from a healthy population (Coriell DNA library (50), n=54) to introduce genetic diversity and increase the number of SNPs tested.

### SNP selection criteria

With the modified design we chose to test 289 PCa risk-associated SNPs which are located on 184 non-overlapping segments that include 50% (70/140) of the previously published PCa risk-associated loci **(Supplementary Table 1; Supplementary Table 2)** (5). These regions were selected as they are potential enhancers which have both H3K27Ac and chromosomal looping to a gene promoter (51). In addition to these risk-associated SNPs we designed the DNA capture to also target 25 strong enhancers (52) (positive control) and 25 inactive regions (negative control) as experimental controls **(Figure 1.A; Supplementary Table 3)**. Using this capture, we enriched randomly fragmented DNA (400-600 bp) that was hybridized with adapters each containing 3bp UMI (53). This was PCR amplified and then cloned into a second-generation STARRseq plasmid. Due to the relatively large size of the insert (median=543bp) it was not feasible to use conventional short-read Illumina paired-end sequencing as variants in these larger inserts potentially will not be identified. Therefore to sequence the entire insert we modified a 500 cycle Illumina sequencing protocol and did two rounds of asymmetric sequencing to include both a “long (450)-short(50)” or “short (50)-long(450)” read **(Figure 1.B)** (43). By matching the unique fragment barcode from the UMI (6bp) and randomly captured insert (18bp) we clustered each pair of asymmetric reads to obtain the full enhancer sequence. Next we collapsed the short and long mates of asymmetric runs and took the consensus sequence (median 36 reads/enhancer; SEM=0.03) **(Figure 1.C)**. In total we reconstructed 32,620 fragments which were located in the capture regions. These asymmetric sequencing results were confirmed with PacBio circular consensus sequencing (CCS) long reads, with 90% (29106/32620) of reconstructed fragments found with Pacbio **(Figure 1.D)**. In addition to the PCa risk-associated SNPs, captured enhancer fragments also included variants from common population SNPs (dbSNP150 common VCF) (9) as well SNPs that were specific to the Coriell DNA population (50). Overall, our SNP library was represented by a median of 67 unique plasmids per SNP containing 41 wild-type (WT) alleles and 26 variant alleles (VAR) respectively. Moreover, each unique plasmid is supported by a minimum of 1000 mapped reads, which is 100x higher than previous work (44). To limit the impact of the potential enhancer position and size bias we calculated the impact of unique plasmids thresholds **(Methods)**. From this we found that having >15 unique plasmids per WT or VAR SNP would minimize the Type-I error to 0.05. Therefore, we focused our analysis on those 308 SNPs that have at least 15 unique plasmids with WT or VAR allele supporting inserts **(Supplementary Figure 1.A; Supplementary Table 4)**. These SNPs include 102 (of 289) PCa risk-associated SNPs (PCa), 194 SNPs commonly found in the 1,000 Genome Project (G5) and 12 Coriell DNA library specific SNPs **(Supplementary Table 5)**. When we compared the variant allele frequency (VAF) with these SNPs we observed a higher but not significant VAF of PCa risk-associated SNPs (*p*=0.56) and G5 SNPs (*p*=0.18) compared to Coriell SNP **(Supplementary Figure 1.B)**.

**Figure 1:**
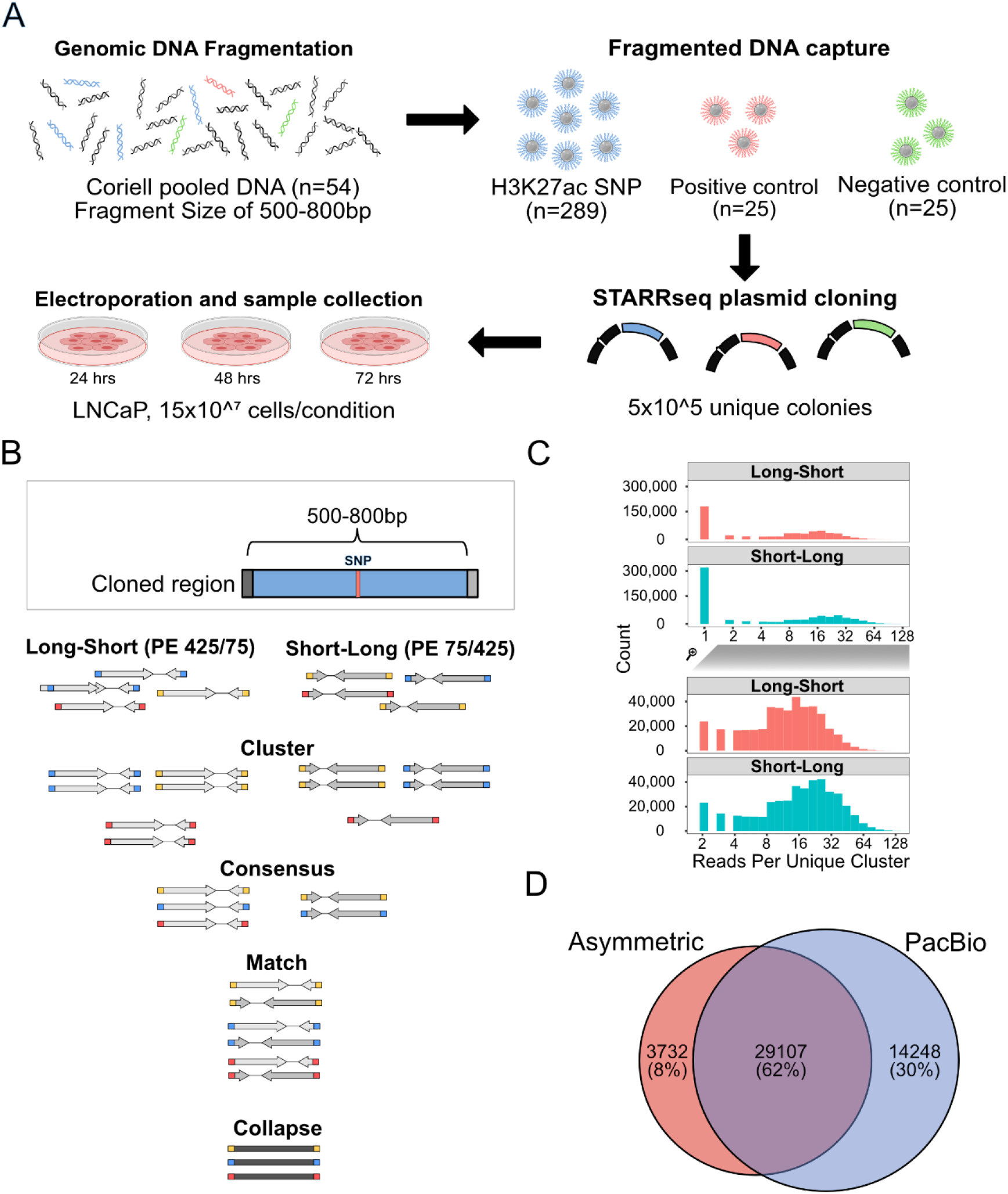
Schematic representation of enhancer fragment sequence reconstruction. a) Experimental steps of snpSTARRseq method. b) Computational analysis of asymmetrical reads. c) Following clustering step, number of read supporting each reconstructed fragment was investigated for each asymmetric run. Top histogram demonstrates the number of single read supported fragments which are removed prior to consensus step whereas the bottom histogram represents those reads included in the further analysis. d) DNA input library was sequenced by Pacbio CCS sequencing to validate presence of reconstructed sequences

### Characterisation of genetic variants that impact enhancer activity

To test the impact of PCa-associated SNPs on enhancer activity, we electroporated the snpSTARRseq library into LNCaP, an androgen receptor (AR)-dependent prostate cancer cell line, and harvested at 24, 48 and 72 hours (n=3 biological replicas). Using self-transcription as a surrogate for enhancer activity we measured the activity of all high-confidence WT and VAR fragments with STARRseq. Following normalization to the input library we observed a clear increase in self-transcription at known enhancers (positive control) but not negative controls with a high correlation of allelic enhancer activity across each experimental time point **(Figure 2.A; Supplementary Figure 1.C).** Principal component analysis (PCA) of all samples demonstrated that 48 and 72 hour samples had a closer enhancer profile compared to the 24 hour samples **(Supplementary Figure 2.A)**. We next investigated how each SNP affected enhancers activity. Using a differential allelic enhancer activity test based on a negative binomial regression model (NBR) we identified 78 unique SNPs across the three timepoints that showed bi-allelic activity at nominal significance level (p value < 0.05) (60, 63 and 43 at 24h, 48h and 72h respectively) **(Figure 2.B; Supplementary Table 6)**. Of these a total of 31 (39%) nominally significant SNPs passed multiple hypothesis testing correction (FDR <0.05) with 23 being PCa disease-associated SNPs identified from GWAS. Further, the majority of the SNPs that impacted enhancer activity (36/78) were PCa risk-associated SNPs, demonstrating the importance of variants on activity. Supporting our PCA analysis, we observed higher correlation in bi-allelic activity across all significant SNPs in the 48 hr and 72 hr samples (Pearson = 0.94) **(Supplementary Figure 2B)**. Focusing on the 48hr samples we observed a similar frequency of activating (n=31) and repressive (n=32) events.

**Figure 2:**
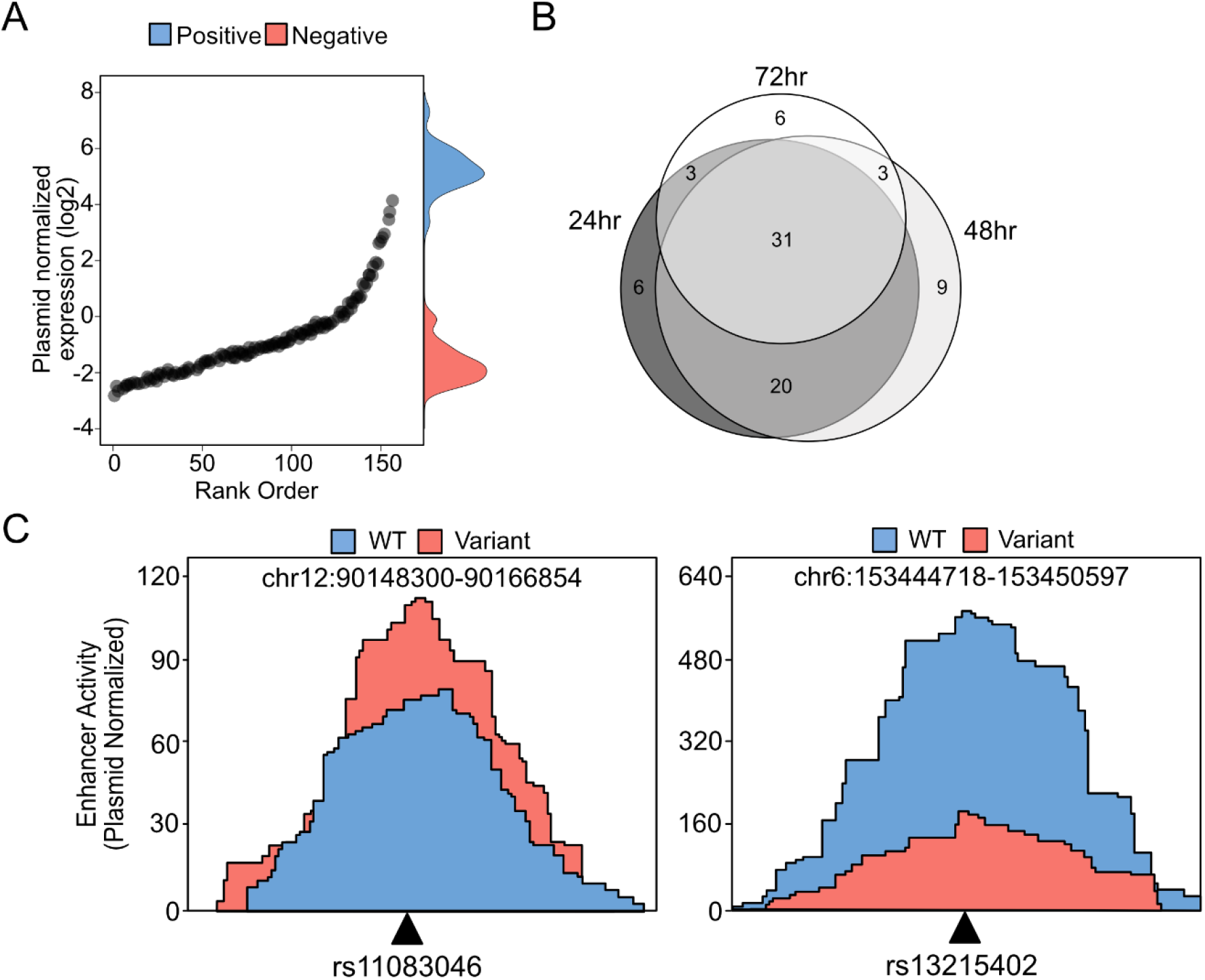
Characterisation of genetic variants that impact enhancer activity. a) Quality control analysis of 48 hour time point demonstrates the SNP, positive control and negative control capture regions normalized count (mRNA/input DNA) distribution. Each black dot represents a single SNP capture region, whereas density plot on the right hand side represents the control capture regions (blue = Positive Control, red= Negative Control). b) NBR model was used to determine SNPs with significant bi-allelic activity. As a result of the calculation, 78 unique SNPs were found among all time-points. Number of overlapping SNPs from each time point is depicted by Venn diagram. c) Activating (rs11083046) and repressive (rs13215402) SNPs that cause bi-allelic enhancer activity were visualized.

Focusing on specific PCa-associated SNPs we observed that rs11083046 (chr18:51781019) variant C allele had a 30% increased enhancer activity (p value = 0.00367) than the reference T allele **(Figure 2.C)**. In contrast, the enhancer activity of rs13215402 (chr6:153447550) decreased by 50% (p value = 0.00349) when the reference allele G was substituted with the variant allele A **(Figure 2.C)**. Supporting these plasmid-based results, those SNPs with bi-allelic enhancer activity commonly affected target gene expression. Using previously published enhancer-promoter interactions from H3K27Ac-HiChIP (51), we found that 20 of 36 “hit” PCa risk-associated SNPs correlated with altered expression of the target gene **(Supplementary Table 7)**. Interestingly, we found 2 significant SNPs (rs13265330; 2.2-fold, FDR=9.25×10^−6^, rs11782388; 1.47-fold FDR=0.07) that were located ~10kb downstream of *NKX3-1,* a gene involved in early stages of prostate tumorigenesis (54). While not significant, we also found supporting evidence of two risk loci enhancers that loop to the PCa-associated genes *CTBP2* and *PCAT19*. Similar to published work two SNPs (rs11672691 and rs887391) near PCAT19 caused increased enhancer activity in all of the time points (14). At the CTBP2 loci, rs4962416 and rs12769019 caused repression and activation respectively similar with previous work (19). Taken together, these results demonstrate that snpSTARRseq can robustly identify those SNPs which alter enhancer activity. When combined with chromosomal confirmation data these results provide a mechanism of action for non-coding disease-associated SNPs.

### Comparison of snpSTARRseq to *in silico* methodologies

*In silico* based pathogenicity predictions are commonly used to stratify non-coding variants for functional characterisation studies. To benchmark the performance of these methods to snpSTARRseq, we obtained the variant impact scores at the tested PCa disease-associated SNPs from ncER (23), CADD (24) and deltaSVM (26). When comparing these *in silico* predictions to our experimental snpSTARRseq results we observed no significant correlation between the deleteriousness (CADD), essentiality (ncER) and deltaSVM score with either bi-allelic effect (log2(VAR/WT)) or statistical significance (FDR) **(Figure 3.A, Supplementary Figure 3.B)**. Further, when we separated SNPs into binary groups based on *in silico* annotations there were no significant changes in the enhancer activity. **(Supplementary Figure 3.C)**. Consistent with the literature (28, 29) we found that *in silico* methods may not provide accurate results to predict enhancer activity.

**Figure 3:**
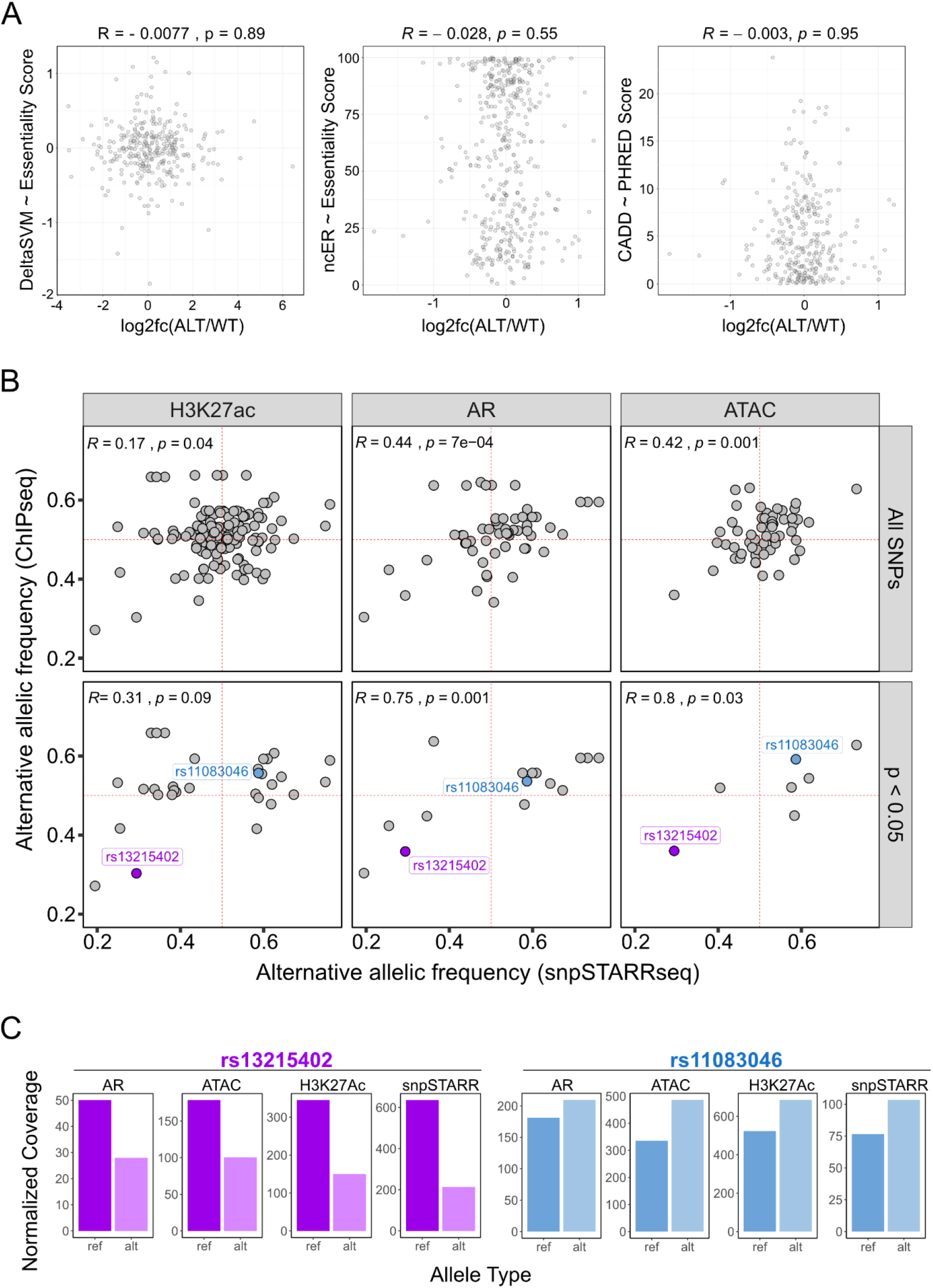
Comparison of snpSTARRseq to orthogonal methodologies. a) snpSTARRseq allelic-effects (VAR/WT; Method; Log2FoldChange) was compared to DeltaSVM (Essentiality Score), ncER (Essentiality Score) and CADD (PHRED score) and no significant relation found. b) snpSTARRseq allele frequency (AF) values were compared against in-vivo AF obtained by H3k27Ac and AR ChIPseq and ATACseq. Top row contains all SNP values without any filtration whereas the bottom row has only significant SNPs found by snpSTARRSeq (nominal p value < 0.05). Two anecdotal examples of repressive (purple) and activating (blue) SNPs were colored to which were found by all 4 methodologies. c) SNPs with activating and repressive effects found by previous *in vivo* work and snpSTARRseq were demonstrated. snpSTARRseq captures the bi-allelic effect of these SNPs accurately.

### snpSTARRseq correlates with clinical allelic-imbalance

We next compared our experimental snpSTARRseq results with *in vivo* allelic-imbalance from H3K27Ac (n=200) and AR (n=131) ChIPseq (33), as well as the assay for transposase-accessible chromatin using sequencing (ATACseq) (n= 26) in prostate tumors (55). Of the 308 SNPs tested by snpSTARRseq, H3K27Ac ChIPseq had the highest coverage among other methods and captured 50% (148/308) of all SNPs. AR ChIPseq and ATACseq both only captured 18% (57/308), of all SNPs tested. Based on our initial comparison without filtration, we observed low to moderate correlation between samples (Pearson; H3K27Ac = 0.17, AR = 0.44, ATACseq = 0.42) **(Figure 3.B, top; Supplementary Table 8)**. However, when non-significant SNPs were filtered out we observed a marked increase in correlation among all comparisons (Pearson; H3K27Ac = 0.31, AR = 0.75, ATACseq = 0.80) **(Figure 3.B, bottom; Supplementary Table 8)**. Based on 48 hr sample, 33% (26/78) of significant bi-allelic SNPs were supported by one, 24% (19/78) by two and 2% (2/78) were supported by all of the allelic-imbalance *in vivo* methods. Those two SNPs that were captured by all methodologies (rs13215402 and rs11083046) demonstrated parallel repression **(Figure 3.C, left)** or activation **(Figure 3.C, right)** of enhancer activity and allelic imbalance. Overall the *in vitro* snpSTARRseq results broadly correlate with *in vivo* allelic imbalance but not *in silico* predictions, highlighting the need for experimental validation of non-coding variants.

## Discussion

The impact of genetic variants on protein-coding amino acid sequence is generally well understood. However, the diversity of activity greatly limits large-scale testing, as each protein requires a specialized assay. Paradoxically, while non-coding variants are poorly understood, the common activity of enhancer CREs makes them extremely amenable to high-throughput screening. In a single experiment thousands of non-coding variants can be functionally tested. Further, when combined with chromosome conformation capture methods these massively multi-parallel assays offer a promising approach to systematically characterize the mechanism of disease-associated non-coding SNPs. However, previous adaptations were either designed to identify new variants (42, 43) or were prone to false-positives (44). Therefore in this work we developed snpSTARRseq to functionally test non-coding genetic variants. Our design utilizes a second-generation STARRseq plasmid, which provides significantly reduced background signal as compared to earlier reporter assays (45). Further, by using a capture-based enrichment of specific genomic loci, we significantly increased the number of target fragments per genetic variant. This is particularly important as the unique number of plasmids can strongly influence Type-I error due to both position and size biases (**Supplementary Figure 1.A**). SNPs tested by our method had a minimum of 30 (WT+VAR) unique plasmids supported by >1000 mRNA reads. This is 100x higher than previous work and dramatically reduces the overall noise caused by variable insert size (44). Next, we tested a larger fragment (400-600 bp) given that enhancer activity generally increases with size potentially due to the additional co-regulatory proteins and TFs that bind to these CREs (56). Testing only small inserts (<250 bp) can be prone to false negatives due to the low signal-to-noise (43). While larger fragments (>600 bp) increase signal strength, they are difficult to sequence with standard short-read Illumina sequencing. We therefore utilized an unconventional asymmetric sequencing methodology to reconstruct larger inserts (500-1000 bp) that were validated with long-read Pacbo. Given the strong consensus, either sequencing approach can be used to identify variants in the plasmid libraries (57, 58). Finally, by using DNA from numerous individuals (>50), we dramatically increased the genetic diversity and number of SNPs tested compared to studies from a single cell line (45). Overall these modifications provide a robust platform for functional testing of GWAS hits.

With this approach we targeted the bi-allelic enhancer activities of 289 PCa risk-associated SNPs from 184 non-overlapping regions that contain a H3K27Ac mark and looped to a gene promoter. Using a very conservative threshold of WT+VAR plasmids, we covered 35% (102/289) of PCa risk-associated variants as well as additional 206 SNPs in the same loci. We observed that bi-alellic SNPs were highly concordant across multiple time points with minor differences between the earlier (24hr) and later (48 and 72 hr) time points. Potentially, this may be due to cells reaching equilibrium in the later time points between snpSTARRseq mRNA transcription and degradation. From those PCa risk-associated SNPs we observed that 35% (36/102) had significantly altered enhancer activity. Many of these significant SNPs were located in 26 previously published PCa risk associated loci (5). Moreover, 72% (26/36) of those with altered enhancer activity were associated with previously published expressive quantitative trait loci (eQTL) (**Supplementary Table 7**). Consistent with the literature, we found supporting evidence for previously published SNPs that alter enhancer activity and impact on expression of *NKX3-1* (rs13215045 and rs11782388) (59–61), *CTBP2* (rs4962416 and rs12769019), *PCAT19* (rs11672691 and rs887391) and RSG17 (rs13215045, 6p25 RSG17 intron variant) (62).

Next, we compared the performance of snpSTARRseq to multiple *in silico* methodologies. Matching previous studies (28, 29), our experimental results did not strongly correlate with any *in silico* method. These differences could potentially be attributed to overfitting and methodological differences (27, 63, 64). Regardless of the cause, the low correlation between different *in silico* techniques (~1%) suggests that there is a need for improved accuracy with these approaches (63). Recent studies have utilized semi-supervised methods to improve results by calibrating calculations and generating cell-type specific predictions (65). Incorporation of single-nucleotide polymorphism evaluation by systematic evolution of ligands by exponential enrichment (SNP-SELEX) experimental results has also been shown to improve model accuracy (66). Contrasting these *in silico* predictions, we observed significant correlation between snpSTARRseq and clinical allelic imbalance of ATACseq, H3K27Ac and AR ChIPseq from tumor tissues. These results support experimental validation of non-coding variants.

To improve the sensitivity and accuracy of large-scale non-coding enhancer assays we developed snpSTARRseq. By increasing fragment length and reducing signal to error this approach can precisely identify functional variants. While not the goal of this work, genetic perturbations of snpSTARRseq systems could be used to identify how specific transcription factor binding is altered by SNPs. Further while focused on germline genetic variants, this same approach can be adopted to study somatic variants. Overall, snpSTARRseq offers an integrated experimental and computational approach to test the bi-allelic activity of hundreds to thousands of genetic variants in a single experiment.

## Data and Code availability

We deposited the snpSTARRSeq computational framework at Github (https://github.com/mortunco/snp-starrseq). All visualization parameters and scripts could be found in publication-figures.rmd file in our repository. All datasets generated during this study along with other processed files are available at SRA under accession PRJNA791664. Supplementary table can be accessed from https://docs.google.com/spreadsheets/d/1IdoTRaXX6Q-Kwk2GI0EG86aknlEK_yxkACsGiycQwSc/edit?usp=sharing

## Acknowledgements

We would like to thank Baraa Orabi and Fatih Karaoglanoglu for their contribution on implementation of Calib and Snakemake respectively.

## Methods

### STARRseq capture library design and additional SNP expansion

We picked 289 SNPs (**Supplementary Table 2**) which are located in enhancer regions that have H3K27Ac signal and chromosomal looping to a gene promoter (51) as well as 50 control regions (25 positive and 25 negative control). All capture regions can be found in **(Supplementary Table 3)**.

### Generation of SNP STARRseq library

Pooled human genomic DNA (NA13421;27 male and 27 female; Coriell Institute for Medical Research; (50)) was fragmented (500-800 bp), end-repaired and ligated with xGen stubby adaptors (IDT) containing random i7 3bp UMI. The target regions **(Supplementary Table 3)** were captured using a custom xGen biotinylated oligonucleotide probe pool (IDT) (53) and Dynabeads M-270 Streptavidin beads (IDT). Post-capture was PCR-amplified with STARR_in-fusion_F primer (TAGAGCATGCACCGGACACTCTTTCCCTACACGACGCTCTTCCGATCT) and STARR_in-fusion_R primer (GGCCGAATTCGTCGAGTGACTGGAGTTCAGACGTGTGCTCTTCCGATCT), and then cloned into AgeI-HF (NEB) and SalI-HF (NEB) digested hSTARR-ORI plasmid (Addgene plasmid #99296) with NEBuilder HiFi DNA Assembly Master Mix (NEB). The SNP STARRseq capture library was then transformed into MegaX DH10B T1R electrocompetent cells (Invitrogen) and the plasmid DNA was extracted using the Qiagen Plasmid Maxi Kit.

### Experimental method for snpSTARRseq

The cloned SNP STARRseq library (100ug plasmid DNA/replica) was transiently transfected into LNCaP cells (5 × 10^7^ cells/replica; 3 biological replicas) using the Neon Transfection System (Invitrogen). Cells were grown in RPMI 1640 medium supplemented with 10% FBS and collected 48hrs post electroporation. These cells were lysed with Precellys CKMix Tissue Homogenizing Kit (Bertin Technologies) and total RNA was extracted using Qiagen RNeasy Maxi Kit (Qiagen). mRNA was isolated with Oligo (dT)25 Dynabeads (Thermo Fisher) and reverse transcription was done with the plasmid-specific primer (CTCATCAATGTATCTTATCATGTCTG). The synthesized SNP STARRseq cDNA was treated with RNaseA and amplified by a junction PCR (15 cycles) with the RNA_jPCR_f primer (TCGTGAGGCACTGGGCAG*G*T*G*T*C) and the jPCR_r primer (CTTATCATGTCTGCTCGA*A*G*C). The SNP STARRseq capture library was PCR-amplified with DNA-specific junction PCR primer (DNA_jPCR_f, CCTTTCTCTCCACAGGT*G*T*C) and jPCR_r primer. After purification with Ampure XP beads, both the SNP STARRseq samples were PCR-amplified with TruSeq dual indexing primers (Illumina) to generate Illumina compatible libraries. RNA samples were sequenced with a HiSeq4000 (150bp; PE) while the DNA STARRseq capture library was asymmetrically sequenced twice with Illumina MiSeq (75/425 PE for first MiSeq run, and 425/75 PE for second MiSeq run).

### Pacbio long-read sequencing

snpSTARRseq input DNA library was digested with NotI-HF (NEB) enzyme for linearization of the plasmid DNA. After that sequence library was sequenced by PacBio SMRT link. Raw Pacbio sequencing data (subreads.bam) was processed by CCS script of SMRT tools (version 9.03) (67). Output CCS file was processed by our pipeline (see section below) to validate reconstructed enhancer fragments.

### snpSTARRSeq computational framework

We developed our computational framework to reconstruct enhancer sequencing using asymmetrical Illumina sequencing data. We described all major steps of analysis methodology in the following sections.

### Asymmetric read clustering

We sequenced snpSTARRseq DNA library with two independent MiSeq runs (“long-short” and “short-long”) with 6.2 million long-short and 9.2 million reads short-long respectively. Prior to clustering, each asymmetric sequencing run was trimmed down to 75 bp fragments from the beginning of the read to match “long” and “short” read lengths. In addition, 3 bp UMI sequences were added to each side of the enhancer fragment to provide diversity. Finally, we used Calib (version 3.4) that incorporates UMI+sequence content of the reads during clustering. Basically, Calib first separates fragments based on UMIs. Then, using minimizer sequences of sequence fragments, it determines the membership of the same or different clusters. We ran Calib with the following parameters `“−e 0 −k 5 −m 6 −l1 6 −l2 6 −t 2 −c 4”` for each independent asymmetric sample. At the end of this step, our reads supporting the same enhancer fragment were grouped together. Calib’s clustering module parameters can be changed upon special cases. Please see for more detailed information (https://github.com/vpc-ccg/calib).

### Consensus read generation of each random enhancer fragment

Calib clustered those reads that belong to identical enhancer fragments into same groups. Next, we use Calib’s consensus module to generate a high confidence sequence of enhancer fragments of each cluster. Clusters with less than <2 reads would be discarded in this step due to lack of support. During the consensus step, Calib builds multiple sequence alignment of members of each cluster. Then determine each base column-wise. Those bases with low consensus are denoted with N. For stricter settings in consensus generation, users can increase the minimum number of reads required from each cluster.

### Matching reads & reconstruct enhancer sequences

Consensus step generates high confidence sequences of enhancer fragments. Until this step, both of the asymmetric samples were analyzed independent from each other. In this step, we merged “long” reads of the asymmetric reads. To do this, we took the first and last 12 bp of each consensus fragment and used it as a fragment barcode. Using fragment barcodes, we matched enhancer fragments from independent samples with each other. There could be two possible outcomes in this step, either an enhancer is supported by both runs, which is paired, or enhancer is only supported by a single sample, which is orphan. We continue the next step with paired enhancers. Given we successfully matched enhancer fragments from both runs, we didn’t need “short” mates anymore. Our asymmetric design leaves minimum 15 bp from both ends of the long reads overlapping. Using this information, we collapsed “long” reads belonging to the same enhancer and retrieved the full length enhancer sequence. For read collapsing, we used the BBmerge (68) software with default settings.

### Creation of variant database

Consensus and read collapsing steps eliminated random sequencing errors on reconstructed enhancer sequences. Next, we aligned enhancer sequences to the reference genome (hg19/grch37) to identify SNPs on the fragments. Each identified SNP was assigned to an enhancer fragment index (fragment startpos:UMI). Then for each different allele, we created a list of fragment indexes supporting WT or VAR alleles. We filtered all of the SNPs with less than 15 unique enhancer fragment support for WT and VAR enhancer fragments. After this filtration, remaining SNPs were annotated with dbSNP150 common VCF file and included into study (9). We deposited 308 high confidence SNPs in the table **(Supplementary Table 4)**.

### Quantification of mRNA data

Until this section, we described individuals steps of the enhancer fragment reconstruction. Remaining steps of the computational pipeline are responsible from the quantification of enhancer activity. snpSTARRseq library transfected LNCaP cells were harvested after 24, 48 and 72 hours. After that mRNA extraction was done as previously explained. Each sample was then sequenced with symmetric 150 bp paired end reads. After that we generated a fragment index for each mRNA read (fragment startpos:UMI). We combined the counts of each fragment in a count matrix which consists of fragment indexes as rows and samples as columns. Given we already obtained a full enhancer sequence, now we can quantify the activity of each enhancer with simply counting enhancer indexes in mRNA samples. What is more, we can quantify the enhancer activity of different alleles using the database we built in the previous step.

### Testing Bi-allelic activity

To identify SNPs with allelic-specific enhancer activity, we conducted a Negative-Binomial Regression analysis to compare the expression of variant-carrying fragments with wildtype fragments. Fragments overlapping at a SNP position were assigned as variant or wildtype based on the allele they carry. Only those SNPs with more than 15 unique plasmids for both variant and wild type alleles were included for analysis. For each SNP, a negative Binomial regression was performed with the following model by using *glm.nb()* function in MASS R library [3]:

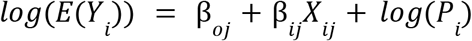

Where *Y_i_* is the RNA read counts of fragment i, *X_ij_* is the allele type of SNP j carried by fragment i, where *X_ij_* =0 when the allele on fragment i is wild type and *X_ij_* =1 when the allele on fragment i is variant type, *β_0j_* is the log expression per plasmid of wild type allele and *β_1j_* is the log fold change of expression per plasmid comparing variant type allele versus wild type allele, *P_i_* is the plasmid DNA read count of barcode serving as an offset term.

### Empirical type-I error for NBR

As the fragment enhancer activity can be affected by not only the variants they carry, but also the specific genomic region they cover, therefore, enough coverage of the SNP to be tested is required to reduce the impact of the position bias. We conducted an empirical analysis to investigate the relationship between number of fragments and FDR. First, we select 10 independent SNPs randomly with at least 30 fragments for each of the VAR and REF alleles, absolute alternative allele effect smaller than 0.1 and p-value larger than 0.5 to treat them as true null SNPs. For each SNP, we downsample the fragments of each allele type to N (where N = 5, 10, 15, 20, 25, 30) and conduct the NBR to test the allelic specific enhancer activity. We repeat the process for 100 times to compute the proportion of tests with p-value < 0.05 as the empirical type-I error at significance level 0.05.

### Asymmetrical sequencing reconstructed fragment coverage distribution

We reconstructed enhancer fragments independently for each asymmetric run as previously mentioned. To investigate the coverage distribution of each reconstructed fragment, we used Calib’s cluster information to visualize coverage distribution and calculated number of reads per cluster. Then used ggplot2’s *geom_histogram* function to visualize distribution of counts.

### Comparison of reconstructed fragments (asymmetric reads) with Pacbio long reads

We compared our fragment library diversity with respect to Pacbio high confidence CCS reads. To do that, we first ran our pipeline separately for Pacbio and asymmetrical Illumina outputs to obtain or reconstruct enhancer sequences respectively. For asymmetrical Illumina, the output file would be “asym.collapsed-fragments.bam”, whereas for Pacbio the file would be “pb.collapsed-fragments.bam”. We filter all unmapped and low quality reads (<MAPQ10). Then process each file with “create-umi-single.py”. This file creates two columns such as “number of eventID observed” and “eventID”. We considered only following chromosomes for further analysis chr1-22+X+Y+Mt. Lastly, we used R’s eulerr (version 6.6.1) (69) package to generate a Venn diagram for the comparison.

### Comparison of all time points with waterfall plot

Using symmetric reads, we counted overlapping DNA and mRNA counts for each capture region for each sample to create a *capture region ~ sample* count matrix. Then annotated capture regions into SNP, positive control (pos) and negative control (neg) groups. Afterwards we calculated the mean value of the capture region coverages for each time point (24, 48 and 72). Finally, we calculated DNA normalized enhancer activity for each capture region. This was calculated by dividing average mRNA coverage (all replicates) to DNA input coverage.

### Principal Component Analysis of all samples

Using the capture *region ~ sample* count matrix, we conducted PCA with R’s *prcomp* function. We further visualized output with ggplot2’s *geom_point* function.

### Comparison with *in silico* methods

The resulting predictions were described as essentiality with ncER (23), deleteriousness by CADD (70), deltaSVM score (change in the chromatin accessibility) by deltaSVM (26). We obtained three different prediction scores tables for each method. For CADD, we created an input file that has 5 columns namely, CHROM, POS, ID, REF, ALT each of our 308 SNPs (CADD website is strict about keeping the header in the input file and keeping the ID column with a dummy variable despite). After that this table was submitted to CADD web browser for obtaining CADD scores (71). For deltaSVM, we created input file that contains “eventID, 10bp flanking reference sequence, 10bp flanking variant sequence” for each SNP. For instance, eventID, chr1;10270386;G;C would have CCGCACCCCTgGTTGAGCCGG as reference and CCGCACCCCTcGTTGAGCCGG as variant sequence (lower capital indicates the WT/VAR SNP allele). Then obtained impact score for each of our SNPs using score_snp_seq.py on deltasvm_models_rme.tar.gz file belongs to predictions on DHT treated LNCaP cell line from Roadmap enhancer models (313) located at https://www.beerlab.org/deltasvm_models (file used: DHS_E2_73_300_noproms_nc30_hg38_top10k_vs_neg1x_avg_weights.out). For ncER, we obtained whole genome ncER predictions(ncER_10bpBins_allChr_coordSorted.txt.gz) from https://github.com/TelentiLab/ncER_datasets and intersected SNP positions using 308 SNP positions **(Supplementary Table 4)** file. After obtaining all of the scores from aforementioned methods, we created boxplots and dotplots using R version 3.6.0 and the ggplot2 package.

### Comparison with *in-vivo* methods

We obtained pre-processed allelic imbalance datasets from three different studies such as H3K27Ac ChIPseq (n=200) (33), AR ChIPseq, (n=131) (33), or ATACseq (n= 26) (55). We matched all datasets (H3K27Ac and AR ChIPseq, ATACseq and snpSTARRseq) using rsID therefore, we dropped 12 Coriell DNA SNPs which did not have corresponding rsID in dbsnp150 common VCF. We shared corresponding AF for snpSTARRseq and *in vivo* methods as well as significance annotation in **(Supplementary Table 8).**

### PCa risk loci analysis

We extracted 147 index SNPs from (5) manuscript’s supplementary table 7 (european SNPs only) and Supplementary table 8. Using rsID(rsXXX), we extracted SNP positions from dbsnp150 common VCF file (hg19). Later intersected with 289 custom capture probe locations. We accepted all interactions within 150 kb distance.

### SNP and gene overlap analysis

We obtained HiChIP-H3k27Ac chromosome interaction data paired-end BED file (BEDPE) from previous work (51). Each line of this file contains one interaction consisting of anchorA and anchorB. Using our in-house script we split anchors and transformed this data into BED format. Then intersected this BED file with hg19/grch37 protein coding gene position (Ensemble v105) obtained from BioMart (72) by bedtools *intersect* function as follows ‘bedtools intersect −a achor.BED −b genepos.bed −wa −wb’. This intermediate file contains anchor-gene information. Similar to previous comparison, anchor BED file then intersected by 308 SNP BED file **(Supplementary Table 4)**. We obtained eQTL information from *Supplementary Table 6* from (51) work. Specifically, those genes that are TRUE for both eGene_TCGA and eGene_THIB columns. Finally, we processed each of these information in SNPs~Gene section of publication-figures.rmd in our github page.

---

**Supplementary Figure 1:**
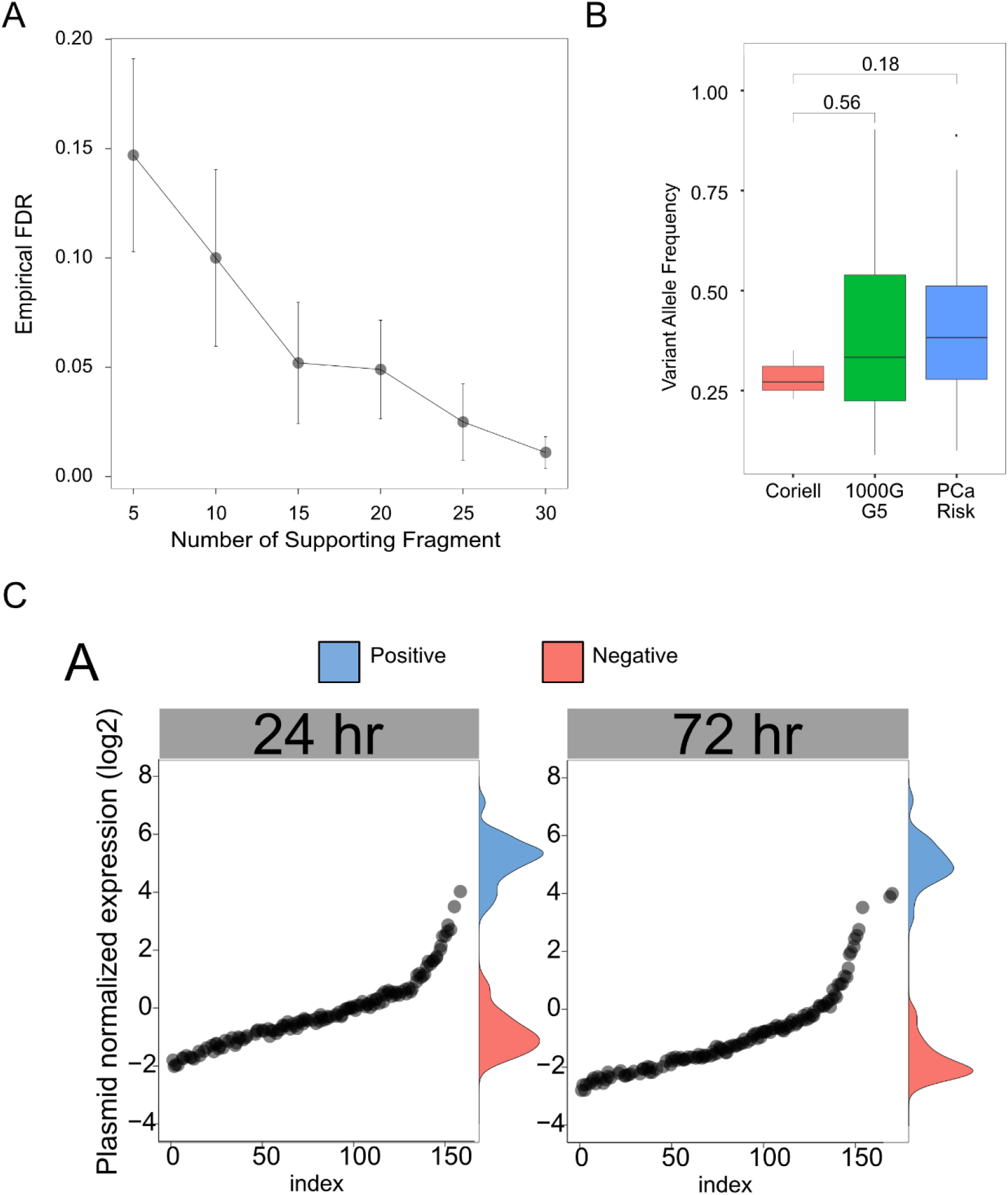
a) Power analysis demonstrates the relationship between number of unique plasmid per SNP and type-I error rate. b) VAF comparison of PCa risk-associated, 1000 genome common and Coriell dataset specific SNPs. c) Quality control analysis results for 24 and 72 hr.

**Supplementary Figure 2:**
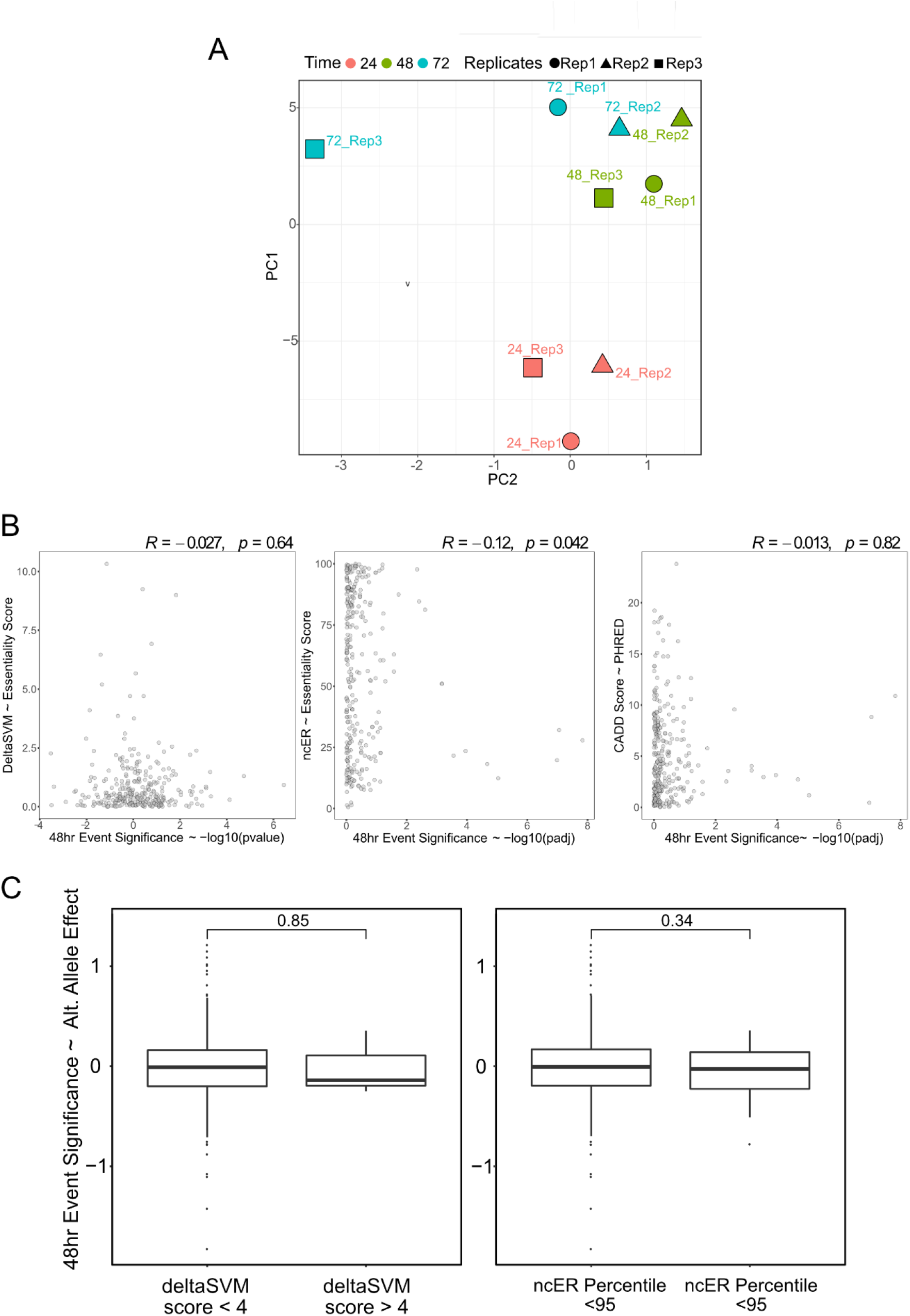
a) PCA analysis of all replicates of 24, 48 and 72 hr samples. b) Allelic-effects (VAR/WT; Method; Log2FoldChange) of SNPs were compared between different time points. d) snpSTARRseq data was analyzed by QuASAR-MRPA and compared to NBR.

